# PARC: ultrafast and accurate clustering of phenotypic data of millions of single cells

**DOI:** 10.1101/765628

**Authors:** Shobana V. Stassen, Dickson M. D. Siu, Kelvin C. M. Lee, Joshua W. K. Ho, Hayden K. H. So, Kevin K. Tsia

## Abstract

**Motivation:** New single-cell technologies continue to fuel the explosive growth in the scale of heterogeneous single-cell data. However, existing computational methods are inadequately scalable to large datasets and therefore cannot uncover the complex cellular heterogeneity.

**Results:** We introduce a highly scalable graph-based clustering algorithm PARC - *phenotyping by accelerated refined community-partitioning –* for ultralarge-scale, high-dimensional single-cell data (> 1 million cells). Using large single cell mass cytometry, RNA-seq and imaging-based biophysical data, we demonstrate that PARC consistently outperforms state-of-the-art clustering algorithms without sub-sampling of cells, including Phenograph, FlowSOM, and Flock, in terms of both speed and ability to robustly detect rare cell populations. For example, PARC can cluster a single cell data set of 1.1M cells within 13 minutes, compared to >2 hours to the next fastest graph-clustering algorithm, Phenograph. Our work presents a scalable algorithm to cope with increasingly large-scale single-cell analysis.

**Availability and Implementation:** https://github.com/ShobiStassen/PARC

## 1. Introduction

Leapfrogging development in single-cell technologies, notably flow cytometry, mass cytometry, high-content imaging cytometry, as well as single-cell RNA-sequencing (scRNA-seq), has revolutionized approaches to measuring a multitude of cellular characteristics (from gene and protein expression, to biophysical and morphological phenotypes) at the single-cell precision. This attribute holds the key not only to defining the diversity of cell types, states and functions, but also understanding how the phenotypic variability within an enormous and heterogeneous population of cells plays a role in tissue development, health, and disease.

Over the years, both the single-cell measurement depth and throughput have drastically been increased and thus have resulted in an explosive growth of large-scale single-cell data. The most prominent is flow cytometry that traditionally offers high-throughput measurements (~10,000 - 100,000 cells/sec), typically with ~10 features (cell-surface markers and intracellular proteins). Integrated with high-speed imaging techniques, imaging flow cytometry has emerged with the simultaneous spurt in measurable parameters and throughput because of its ability to allow high-resolution single-cell image-based phenotyping without compromising the throughput significantly (Caicedo et. al., 2017; Blasi, T. et al. 2016; Lee et. al., April 2019). Mass cytometry by time of flight (CyTOF) offfers single-cell measurements of millions of cells with simultaneous detection of 40 or more proteins (features) for a given experiment (Spitzer and Nolan, 2016), albeit at a lower throughput compared to flow cytometry. Another parallel advance is scRNA-seq, recent years have seen proliferation in scRNA-seq data with droplet-based systems sequencing hundreds of cells per second (Zheng *et al.* 2017) resulting in tens of thousands of cells across samples in an experiment. A “MegaCell Demonstration” by 10X Genomics (10X Genomics Datasets, 2017) recently featured a scRNA-seq data set of 1.3 million E18 mouse brain cells, showcasing the very high throughput it can achieve.

While the single-cell measurement scale or throughput continues to grow at a staggering rate, such technological advance has outstripped the existing computational capability to handle, process and analyse the enormous amount of heterogeneous single-cell data. Beyond the sizeable single-cell data backlog, the key challenge lies in the search for new solutions to fill such a computational gap and thus to accelerate new biological discoveries. Among all computation tasks, unsupervised clustering plays a decisive role in facilitating downstream biological interpretation in single-cell analysis. However, available methods still lack the scalability and data-driven capability required for parsing the immense and heterogeneous data and thus identifying putative cell types in an efficient manner.

Most tools developed for gene sequence data become computationally prohibitive when the cell count reaches 10^5^-10^6^ cells. For example, to handle the scRNA-seq data of only 6,000 cells, the popular SC3 and RaceID take ~20,000 seconds and CIDR takes 1000 seconds (Duo et al. 2018). Even the clustering stage of the speedier Seurat takes over 30 minutes on 60,000 cells (Sinha et al. 2018). In order to digest larger batches of data, the common strategy is to rely on subsampling which inevitably overlooks rare cell types (e.g. SPADE Qiu et al. 2011). A handful of tools can operate on large cytometry data sets, e.g. FlowSOM (Van Gassen et al. 2015), K-Means and FlowMeans (Aghaeepour et al. 2011). However, they often rely on manual parameter tuning, or invoke a number of clusters in advance that in turn poses challenges to perform unbiased exploration of the unknown complex cellular heterogeneity. In the scenario where it is feasible to perform analysis for a range of predetermined number of clusters and select the result based on the internal clustering evaluation criteria (e.g. Silhouette Index), it is not uncommon that the emerging optimum or ‘elbow point’ is a poor reflection of the true underlying structures in the data. A recent benchmarking study of 12 clustering methods on smaller scRNA-seq data sets (Duò et al., 2018) showed that generally no method achieved its best performance at the annotated number of clusters. For instance, in its automated mode where cluster-selection is based on the elbow-point of a performance metric, FlowSOM vastly undershoots the number of clusters (as does FlowPeaks), and typically requires a ‘generous’ cluster estimate in order to capture annotated populations (Weber and Robinson 2016).

In light of these challenges, we present PARC, *phenotyping by accelerated refined community-partitioning* - a fast, automated, combinatorial graph-based clustering approach that integrates hierarchical graph construction and data-driven graph-pruning with a new community-detection algorithm. PARC (i) outperforms existing tools in scalability without resorting to sub-sampling of ultralarge-scale, high-dimensional single-cell data (> 1 million cells); (ii) accelerates the clustering computation by an order of magnitude through community-partitioning refinement; and (iii) more importantly, augments the sensitivity and specificity to unbiasedly recapitulate the cellular heterogeneity, especially rare subsets within large populations. We validate the superior performance of PARC on large-scale datasets, with respect to speed and accuracy, as well as versatility across a wide range of single-cell data including: mass and flow cytometry, scRNA-seq and imaging cytometry. Notably, we demonstrate that PARC not only can detect subpopulations that were not labelled in the original scRNA-seq data sets of 68k peripheral blood mononuclear cells (PBMC), but it can also enable data-driven clustering of the entire mouse brain dataset of 1.3 million cells without any downsampling. As a proof of concept, we also showcase that PARC correctly infers cell type on a megaset of multiple lung cancer cell lines (> 1 million cells) on the basis of their intrinsic, biophysical attributes derived from multi-contrast label-free single-cell images (Lee, Feb 2019 and Lee April 2019).

## 2.1 Method

PARC is built upon a nearest-neighbor graph architecture in which each node is a single cell, connected to a neighborhood of its similar cells by a group of edges (**Fig. 1a**). PARC employs three major innovative steps to enable fast, scalable, and accurate data-driven clustering of heterogeneous single-cell data. PARC first constructs the graph based on an accelerated k-nearest-neighbor (k-NN) search using hierarchical small world (HNSW) (Malkov and Yashunin, 2016). In contrast to the tools adopting exact neighbor searches (e.g. X-Shift (Samusik et al 2016)) whose computational overload is hardly justified in terms of accuracy, HNSW-based k-NN search offers logarithmic complexity scaling. Second, data-driven graph pruning (on both local and global scale) is implemented in order to remove extraneous edges guided by the distribution of edges weighted by the Jaccard and Euclidean metric. This step is motivated by the distribution of weights which generally resembles a long-tailed distribution in various single-cell data sets in consideration (**Fig. 1b**). Based on the histogram we observe that the Jaccard (and also Euclidean) weight for weak (potentially ‘spurious’) and majority (around median weight) links is similar, conceivably a consequence of the curse of dimensionality. The tail thus carries most of the important neighbors, but its high weight score also diminishes the relative difference between weak and majority links. Consequently, the optimization function employed in the subsequent community-detection step sees the weak and majority links as very similar. The detected subcommunities are thus more susceptible to being merged by spurious links due to the “resolution limit” – a common limitation in community detection (Barabasi et. al. 2019) resulting in undesirable merging of clusters. We find that true communities have adequate strong and ‘majority’ level weight neighbors to still emerge as standalone clusters even after pruning (evidenced by no excessive fragmentation of communities in the final output). We also find that pruning considerably improves sensitivity to rare but distinct populations. Hence, the data-driven pruning procedure offers a two-fold impact in reducing the sample size of edges and improving the k-NN graph representation of the underlying data structure, both of which critically improve the subsequent community detection step in speed and robustness. Third, a newly developed community-detection approach, Leiden algorithm (Traag et al., 2019), is employed to partition the large pruned networks in the graph into communities. In contrast to the popular Louvain algorithm (Blondel et al. 2008), Leiden algorithm demonstrates superior performance in faster computation time, scalability, and minimising badly connected communities (Traag et al. 2019).

**Fig. 1.**
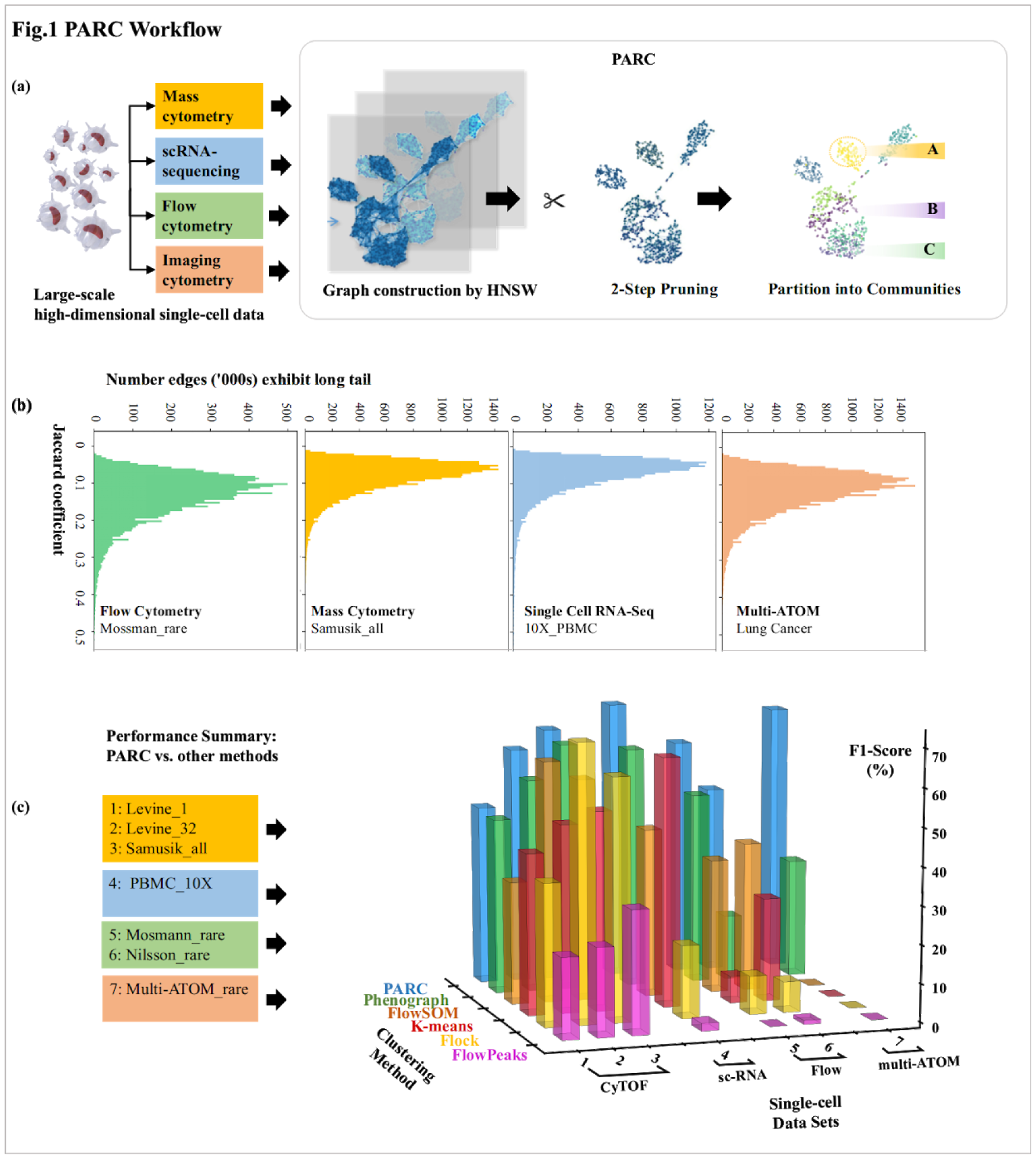
Overall Workflow and Performance of PARC. (a) Overview of PARC workflow for large-scale single-cell analysis on multiple types of high-dimensional single-cell data. The enabling features include fast graph construction by HNSW, 2-step data-driven graph refinement and pruning, and accelerated community detection by Leiden algorithm. (b) Long tailed distributions of graph edge-weights observed in various SC datasets. The tail implicates that the high weight score of the important neighbors diminishes the relative difference between weak and majority links. This negatively impacts the robustness and the speed of community detection – a predicament that could be addressed by graph pruning. (c) Overall performance of PARC and other competitive clustering methods on various SC datasets.

The data input to PARC is a matrix of *n*_*cells*_ x *m*_*dims*_ representing *n*_*cells*_ single cells, each of which has *m*_*dims*_ phenotypes (dimensions) measured by different single-cell technologies, e.g. flow cytometry, CyTOF, scRNA-seq, and imaging cytometry (e.g. multi-ATOM in this study), etc. We next describe the three key modules mentioned above that augment the clustering speed, scalability and accuracy, particularly in the case of ultralarge dataset.

### 2.1 HNSW for fast and scalable K-NN search

In PARC we use HNSW as an approximate nearest neighbor search based on the assumption that our data has an inherently community like structure. We note that some tools (e.g. X-Shift Samusik et al 2016) employ exact neighbor searches whose computational overload cannot be justified in terms of accuracy. Furthermore, the appropriate choice of an approximate nearest neighbor algorithm is essential for scalability. Several others incorporate approximate nearest neighbor searches that become time intensive for use on large-scale data (e.g. Phenograph’s use of Python LibrarySklearn’s ‘kdtree’ and SCANPY’s UMAP (McInnes et al. 2018) based neighbor-search). A small world graph is characterized by long links which bridge different clusters and shorter links which represent inter-cluster connectivity. The HNSW method differs from other NSW methods by binning links in hierarchy (in layers) according to their lengths. The search starts at the top layer containing the longest links, and traverses the elements until a local minimum is reached. The search then goes to the lower layer (i.e. the layer having shorter links) from the node where the most recent local minimum was detected. Such hierarchical graph structure allows fast graph construction with logarithmic scalability, i.e. the construction scales as O(NlogN), whilst each query takes O(logN) time (Malkov and Yashunin, 2016).

### 2.2 Graph pruning for effective capture of network structure

The linkages present in the K-NN graph impact the number of clusters found in the community-detection algorithm (more precisely, modularity optimization). Instead of using the common strategy of tuning the K parameter, PARC starts with a generous fixed K number (K = 30) and implements automated weak-edge pruning guided by the data structure. Not only can this data-driven pruning approach effectively ‘fine-tune’ the local K value, but also refine the network structure for accelerating and optimising the Leiden community detection steps.

A user-defined global K value and its tuning do not yield robust graph representation of the data. Higher K values generally favor preserving larger communities, but compromise the ability to detect rare subpopulations. One might expect, on the other hand, that a lower value can automatically separate distinct rare subpopulations, but leads to more severe cluster fragmentation - complicating the biological discovery. We found that using a graph of a lower k-value as input to the modularity algorithm only marginally (and inconsistently) allows us to discern rare populations compared to a larger K value (See **Fig. 2**). Furthermore, in a heterogeneous single-cell dataset, the degree of each node (cell) often varies across the K-NN graph. The baseline degree of a node also exceeds K, and spurious connections exist. Choosing a global K value thus does not allow that density of links to vary sufficiently. Even worse, as the construction (and querying) of the graph is based on the Euclidean distances in higher dimensional space, the relative importance of neighbors is not necessarily accurately quantified due to the common ‘curse of dimensionality’, and re-weighting the graph based on different metrics to will still suffer the same “curse of dimensionality”.

**Fig. 2.**
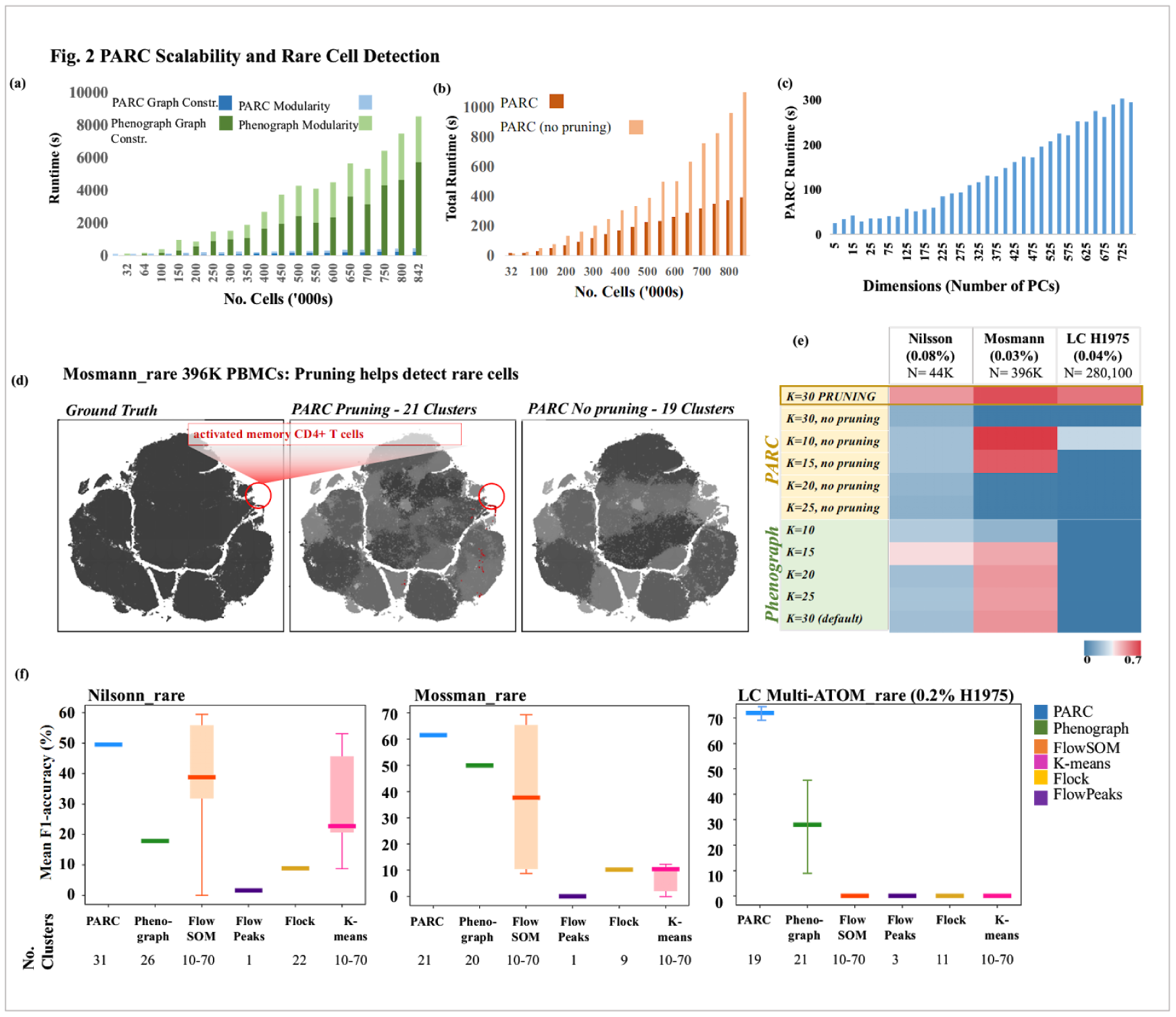
Scalability of PARC and its Rare Population Detection Performance. (a) Scalability of PARC, in comparison to Phenograph, in terms of graph construction and clustering time, on the random samples of large-scale CyTOF data (Samusik_all: 841,644 cells). (b) Pruning speeds up the runtime of PARC by a factor of 2, with the gain being greater as the number of samples increases. (c) PARC scalability as dimensionality increases on scRNA-seq data (10X_PBMC). (d) (Left) t-SNE plot colored by ‘ground truth’ of the Mosmann_rare data ; (Center) t-SNE plot colored by PARC clustering with pruning where the cluster containing majority of rare activated memory T-cells are colored red; (Right) t-SNE plot colored by PARC clustering without pruning. The rare activated memory T-cells are not detectable at all. (e). Pruning is key to reliably identifying rare cell population in 3 datasets with rare populations: Nilsson_rare, Mosmann_rare, multi-ATOM_rare, with the rare populations of 0.08%, 0.03%, and 0.04%, respectively. Simply lowering the K (number of neighbors) in graph construction does not ensure rare-cell detection in PARC or Phenograph. (f) Performance comparison of PARC on 3 rare-cell datasets against 5 competitive tools and their corresponding number of clusters.

As discussed earlier, empirically we observe for the various biological data sets in consideration, that the edge-weight statistics resemble a long-tailed log-normal distribution whose tail diminishes the distinction of more closely separated points. A graph that does not discriminate adequately between links (especially those connected to rare populations) dramatically impacts the modularity optimization in the Leiden algorithm.

Motivated by this characteristic, PARC removes ‘weak’ edges in two steps in order to overcome the aforementioned shortcomings brought by the K-value. First by examining each node locally and removing the weakest neighbors of that particular node based on the Minkowski distance as a generalized distance measure; and second by re-weighting the edges using the Jaccard similarity coefficient and globally removing any edges below the median Jaccard-based edge weight. The Jaccard coefficient is the ratio of the intersection of node A and node B’s neighbors, to the union of node A and node B’s neighbors, expressed as:

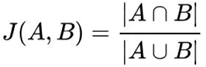

The local pruning allows us to remove redundant neighbors in very densely connected neighborhoods, and the global pruning removes spurious edges that were placed in otherwise disconnected, low-density populations. This results in a graph where only significant neighbors are retained and has the additional effect of making the network even sparser, allowing for a rapid convergence of the optimization algorithm which empirically scales linearly with the number of nodes in a sparse graph (Blondel et al. 2008).

### 2.3 Leiden algorithm on the *pruned* graph to shield against “resolution limit”

Having constructed a network representation of the single cells, i.e a collection of cells that have strong links to each other but fewer links to the remaining network, we apply a new modularity optimization algorithm called Leiden algorithm, for community-detection-based clustering (Traag et al. 2019). Leiden method has very recently been demonstrated to show superior performance in speed as well as reducing internally disconnected (or poorly connected) communities, to the popular Louvain algorithm in clustering social network datasets. Here we apply Leiden algorithm on the biological single-cell data and show that a pruned graph, as an input to this algorithm, could further accelerate the clustering time and mitigate the ‘resolution limit’ issue (whereby smaller clusters tend to become more likely subsumed into larger ones as the network size grows) of the modularity optimization in the algorithm.

A popular method for community detection, modularity optimization is an NP-hard problem, i.e. it is computationally challenging to solve and requires heuristic approximations when applied to large networks. However, the Leiden algorithm stands out in terms of scalability. Modularity is a scalar measure of the density of links within a community to that between communities. For weighted networks, the modularity is defined as (Blondel et al., 2008):

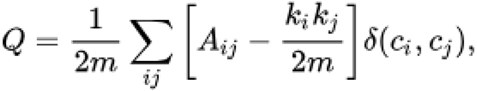

*A*_*ij*_ is the weight of the edge between vertex *i* and *j, k*_*i*_ is the sum of vertex *i*’s edge-weights. *c*_*i*_ is the community to which vertex *i* is assigned. *m* is the weight of the graph. The Kronecker delta function is 1 if *c*_*i*_*=c*_*j*_, and otherwise 0.

Asides from the ‘resolution limit’ issue of the modularity optimization function which tends to merge small sub-structures, the Louvain algorithm applied to any quality function suffers from a separate issue of poorly connected or even internally disconnected communities (Traag et al. 2019). Recall that the Louvain algorithm essentially consists of repeating two steps: 1) re-assigning nodes to communities based on improvement in modularity and 2) once nodes can no longer be moved, aggregate nodes in a community and allow each community to become a ‘node’ in the next pass of the same steps. Leiden addresses the issue of internally disconnected communities by allowing clusters to be refined, broken up into sub-clusters, which can then be assigned to any of the existing major clusters found in the aggregation step immediately before. This vastly improves the connectivity within clusters. However, in order to control the proliferation of clusters Leiden only re-assigns refined communities to those un-refined communities found in the previous step.

This means that once a community is merged into another (which may occur more than desired due to the resolution limit), it can only be reassigned to any of the existing other communities or itself. Thus, even though nodes within communities may be well connected, sub-structures may be subsumed/integrated into larger communities due to the quality function which is sensitive to spurious links extending from minor-populations to major populations. The effect worsens as the size (*m*) of the network increases. This resolution limit clarifies why both Phenograph (which uses Louvain community detection method) and Leiden-without any pruning are unable to consistently segregate rare yet distinct populations. The change in modularity when assigning a node *i* in community *A* to community *B* can be written as (where *k*_*i,in*_ is the sum of the weighted links from node *i* to nodes in *B, k*_*i*_ is the weighted links incident on node *i, Σ*_*tot*_ *i*s the sum of weighted links incident on *B)* :

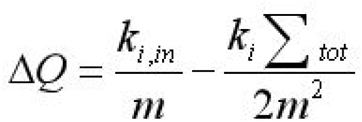

If we were to assign all the nodes in community *A* to community *B* then the change in modularity as a result of merging communities *A* and *B* is

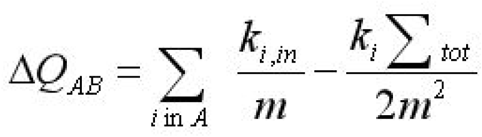

For the simplified case of an unweighted graph (or a graph where the weightings are not discriminatory enough and hence effectively unweighted), we can rewrite the change in modularity when we merge *A* and *B* as, (where *k*_*A*_ and *k*_*B*_ are the total degrees of *A* and *B* and *L* is the total number of links in the entire network, and *l*_*AB*_ is the number of links between community *A* and *B* (Barbasi 2019):

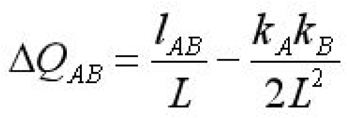

Consider the scenario where *k*_*A*_*k*_*B*_/2*L* <1, then the change in modularity is positive as long as there is even just one link between the two communities (*l*_*AB*_>=1). For the sake of simplicity, let *k*’=*k*_*A*_ ~ *k*_*B*_, then the modularity increase will be positive when A and B are merged for all *k*’≤√(*2L*). So if the number of links within a certain small community are below the threshold √(*2L*), then even a single link to another community will result in a merger and the algorithm will struggle to resolve communities below the resolution limit of *k*’≤√(*2L*). It is therefore critical to remove artificial or weak links set up in the initial K-NN graph.

In some cases, pruning yields clusters consisting of only a single or two cells. In order to determine whether or not these are true outliers, we examine any clusters below a certain minimum population (default of 10 cells). These cells are assigned to the cluster containing the greatest number of its original neighbors found in the HNSW stage, provided this cluster is above the minimum population threshold. If the cell does not have any neighbors belonging to a larger cluster, then it remains an ‘outlier’ cluster. A default value of 10 cells is applied to datasets in this paper.

### 2.3 PARC algorithm

The overall workflow of PARC is summarized in the pseudocode below:

<u>PARC Pseudocode</u>

**Inputs:** X = data [n_cells_ x m_dims_],

global_jaccard_prune = “median”, local_euclidan_prune = “2 standard dev”, small_cluster_threshold = 10 cells

**Outputs**: Cluster membership

*hnsw_graph* <-make_small_world(X) // construct the HNSW graph

*(neighbors, distances)* <-extract_neighbors(*hnsw_graph)* // query 30-NN and weights based on Euclidean distances

**for** each **node_i** in graph // local node pruning

Remove edges 2 standard deviations above mean edge-weights of node_i

// update neighborhood information

*Jaccard-weights* <-compute_Jaccard (on updated edges of graph) // use iGraph library

*Updated edges* <-keep edges above median jaccard-weight of graph

*Simple graph* <-combine edges by summing weights// remove directionality

*Clusters* <-Leiden_modularity_optimization (*Simple Graph*)

**for** each small_cluster in Clusters: // default any cluster below 10 cells

**if** cells in small-cluster have neighbors from original 30-NNG belonging to larger identified clusters: re-assign possible.

**else:** leave as outlying or ‘extremely rare’ population

## 3. Results

With large-scale clustering being the emphasis, we reviewed and compared 18 well-known clustering tools recently benchmarked (Weber and Robinson 2016). Only 5 of them are practically scalable and run to completion on datasets with ~1 million cells without the need for any subsampling (typically subsampling is only applied when runtimes exceed tens of hours or even days). These 5 are Phenograph, FlowSOM, FlowPeaks, Flock and K-Means.

Motivated by the need for a versatile tool to cope with the increasing diversity in the available large-scale single-cell data types, we tested PARC on a range of annotated single-cell data sets of scRNA-seq, flow cytometry, CyTOF, and imaging cytometry with the cell counts spanning almost 2-orders-of-magnitude (from 68,000 to 1,300,000 cells). PARC’s performance is benchmarked against the 5 competitive clustering methods chosen for their ability to also handle large-data sets and a summary of the results is illustrated in **Fig. 1c and Supplementary Fig. 1**. PARC consistently outperforms, in terms of unweighted F1-measure calculated using the Hungarian algorithm (**Supplementary Information**), against Phenograph and other competitive methods such as FlowSOM (Van Gassen et al. 2015) and FLOCK (Qian et al. 2010), especially in revealing minor populations without artificially fragmenting larger populations. In addition, we note that the scores for K-means and FlowSOM shows high variability strongly depending on the pre-determined values of chosen parameters (e.g. K clusters) (**Supplementary Fig. 1**).

In the following sections, we will demonstrate the usability of PARC on three diverse types of single cell data: (1) flow cytometry or mass cytometry data, (2) transcriptomic data, and (3) imaging-based biophysical properties.

### 3.1 PARC is scalable and accurate to very large single cell mass cytometry data

To evaluate systematically how PARC accelerates graph-based clustering, we compare the runtime break-down between PARC and Phenograph in terms of their graph construction and modularity optimization steps. Here we employ the large-scale CyTOF data set Samusik_all: a dataset consisting of replicate-bone-marrow of 10 individual C57BL/6J mice, of 841,644 cells with 13 surface markers (Samusik et al 2016). The initial value of K (the number of nearest neighbouring cells) is 30 for both PARC and Phenograph (which applies the Louvain algorithm). By randomly subsampling the data set with varying size, we quantify how different steps contribute to the computation speed-up in PARC (**Fig. 2a**). Clearly, both the graph construction and modularity optimization are drastically accelerated in PARC, altogether which lead to ~30 times faster in total runtime compared to Phenograph. This speed up may be attributable to some key innovative steps in PARC, namely 1) the use of HNSW to accelerate the nearest neighbor search, and 2) the pruning phase which has a knock-on effect to speed up the modularity optimization by effectively reducing the number of edges for a given number of samples. Notably, pruning progressively becomes more significant in lowering runtime with increasing sample size (**Fig. 2b**) - marking its significance in clustering acceleration with large-scale single-cell data. We also tested the scalability of PARC with increasing data dimensionality using the scRNA-seq dataset of human PBMCs (Zheng et al., 2017). We observe a fairly linear scaling in runtime of PARC, even when the dimension goes beyond 500, indicating its ability to scale with high dimensionality (**Fig 2c**).

### 3.2 PARC identifies rare populations in large flow cytometry data

We next test the ability of PARC to isolate rare populations (See **Fig. 2d-f**) on two flow cytometry datasets available on FlowRepository (repository I.D.: FR-FCM-ZZPH) (Weber and Robinson, 2016), and one in-house imaging flow cytometry dataset of 7 lung-cancer cell lines (representing the 3 major subtypes of lung cancer) generated by a new ultrahigh-throughput label-free imaging technique, coined multi-ATOM (Lee Feb 2019 and Lee April 2019) (See Supplementary Information for experimental details). The first is Nilsson_rare (Nilsson, 2013) which contains 44,100 bone marrow cells with 13 surface markers (dimensions), out of which we must isolate 358 (0.08% of total population) manually gated hematopoietic stem cells. In the second data set, Mosmann_rare (Mosmann, 2014), we have 396,400 human peripheral blood cells (14 surface markers), stimulated with influenza antigens. Only 109 (0.03%) of these are manually gated as activated memory CD4 T cells (see **Fig. 2d**). The third set, *multi-ATOM_rare*, consists of digitally mixed cells obtained from 7 different lung-cancer cell lines with 26 quantitative biophysical features extracted from each label-free single-cell image. Within this dataset, there are only 100 randomly subsampled adenocarcinoma cells (H1975) and 280,000 from the remaining 6 lines (0.04% in **Fig. 2e)**. In these datasets, the cluster with the highest F1-score for any cluster containing members of the rare population is reported (**Fig. 2e and 2f**). The F1-score of the 100 H1975 adenocarcinoma cells are averages across 10 runs with randomly sampled subsets.

PARC consistently outperforms other methods in detecting rare populations across these different datasets (**Fig. 2f**). The F1-score of the rare population obtained using the common large-scale methods (notably FlowSOM and k-means) are not only lower, but sensitive to the user-defined choice of {k =10,15,…60}. Testing on the *Mosmann_rare* dataset, we show that pruning in PARC critically enables the detection of the small activated memory CD4 T cell population (0.03%), which is otherwise missed if pruning is skipped (**Fig. 2d**).

We note that in Phenograph the ability to identify rare but distinct cells is influenced by the choice of number of K (nearest neighbors) in the graph. Although the rationale in Phenograph for applying weights to the edges of the graph is to resolve rare populations by weakening spurious links, we find that the weighted values are not adequately discriminatory as already illustrated in the long-tailed weight distributions. Consequently, a critical factor in faithfully capturing the network structure is whether or not a link exists. A false remedy is to lower K (number of nearest neighbors), however, as shown in **Fig. 2e**, reducing K in PARC or Phenograph’s graph construction does not yield reliable rare-population identification, not to mention the resulting over-fragmentation of clusters that confounds downstream analysis..

The ability of PARC to minimize the risk of over-fragmentation is also evident by comparing it with another clustering method, X-Shift, using density-based K-NN estimation. This method was originally established for analyzing the CyTOF single-cell data “Samusik_all” (Samusik et al 2016). X-shift generated 74 clusters in ~230 minutes based on a random subsampling of 300,000 cells from Samusik_all data, whereas PARC managed to produce 24 clusters of the 841K cells in only ~6 minutes, and to maintain the same F1 scores (**Supplementary Table 1**). Further tested with another CyTOF data sets of healthy human bone marrow cells (Levine_13dim consisting of 24 manually gated populations), this density-based K-NN estimation approach yields over-fragmentation (153 clusters after ~2500 seconds), in stark contrast with PARC (25 clusters after 35 seconds).

### 3.3 PARC dissects heterogeneous single-cell transcriptomic data of PBMCs and 1.3 million mouse brain cells

We also tested the adaptability as well as scalability of PARC in handling complex single-cell transcriptomic (scRNA-seq) data. We use a mid-sized annotated 3’ mRNA data set of 68,000 peripheral blood mononuclear cells (PBMCs) (Zheng et al. 2017) for a more granular analysis, and an exploratory large dataset of 1.3 million embryonic mouse brain without subsampling.

In the first case, the cells in the mixture were annotated by correlating (Spearman) each cell against the average expression profile of 11 purified populations (Zheng et al 2017). We adopt the same pre-processing steps as Zheng et al 2017 which are: by filtering out genes based on unique molecular identifier (UMI) count, selecting the 1000 most variable genes and subsequently using the first 50 principal components (PCs) generated by the principal component analysis (PCA) applied on the UMI counts. Likewise, the issue of ‘drop-outs’ is not directly addressed, but partially mitigated by UMI-count-based filtering. We compute the log2 fold changes by cluster to infer the cell population based on the most differentially expressed genes (**Fig. 3b**).

**Fig. 3.**
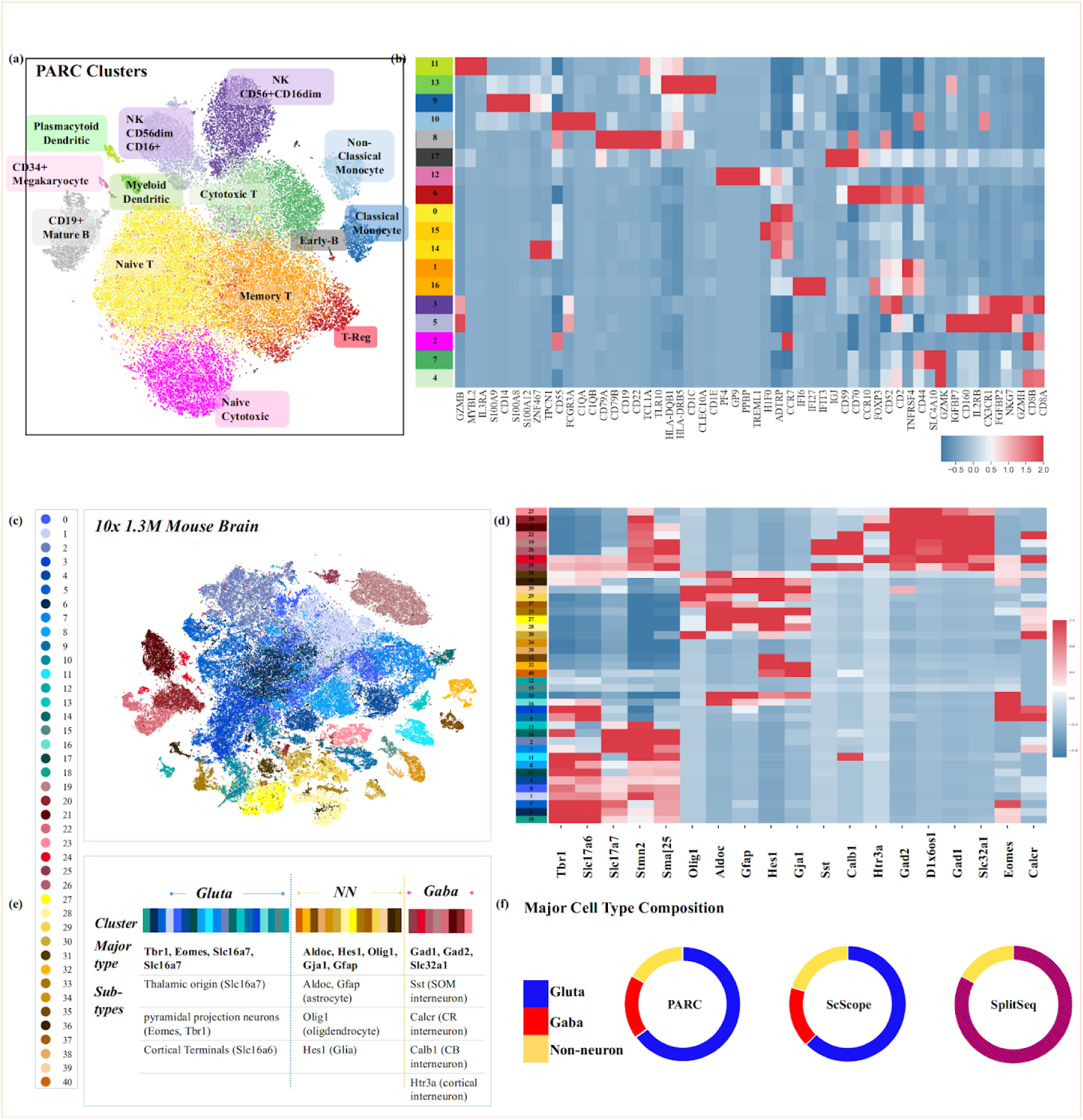
PARC for Mid-scale (68K cells) and Ultralarge-scale (1.3M cells) scRNA-seq Analysis. (a) t-SNE visualization of 68K human PBMCs (Zheng et al. 2017) colored based on clusters found by PARC, which delineates well-known cell subtypes not captured in original annotation (Supp. Fig. 4, Supp. Table 3 for detailed references of marker genes)). (b) Heatmap of most differentially (log2-fold) expressed genes in each assigned cluster by PARC. (c) t-SNE visualization of the entire mouse brain data (1.3M cells). Cluster colors reflect PARC clustering of major neuronal type (Glutamatergic, Gabaergic and non-neuronal) inferred by the marker genes (Allen Brain Atlas and Tasic 2016, Tasic 2018). (d) Mean cluster-level gene expressions of known marker genes and (e) the inferred sub-cell types. (f) PARC’s major cell-type composition concurs with ScScope and SplitSeq, as well as prior studies on embryonic mouse brain cells (See Suppl. Table 4).

Apart from that PARC yields better F1-scores than other clustering methods (**Fig. 1c** and **Supplementary Fig. 1**), it also identifies sub-populations (**Fig.3a**) that were masked by the original manual gating (**Supplementary Fig. 4**). This is attributed to the fact that the annotation was mainly given to T-cell subpopulations on a *mesoscopic* level (e.g. CD4+, CD8+, memory and regulatory T cells). In contrast, other sub-types of PBMCs (e.g. monocytes, dendritic cells and NK cells) are not annotated by any of their known subtypes. Nevertheless, PARC is able to reveal the clusters showing high expression of CD14 (cluster 9) and CD16 (or FCGR3A) (cluster 10), markers for classical and non-classical monocytes respectively (Ong, 2018). It also identifies subsets of NK cells as inferred by the expression level of CD160 and CD16 (FCGR3A) (cluster 3 and 5), which is known to be associated to the CD56dim CD16+ cytotoxic NK cell phenotype (cluster 5) (Bouteiller, 2011). Notably, PARC also detects rare populations of IL-3RA+ (Zhang et al., 2017) plasmacytoid dendritic cells (cluster 11, 0.6%) and megakaryocytes (cluster 12, 0.4%). The marker genes identified for each cluster are summarized in **Supplementary Table 3**.

**Fig. 4.**
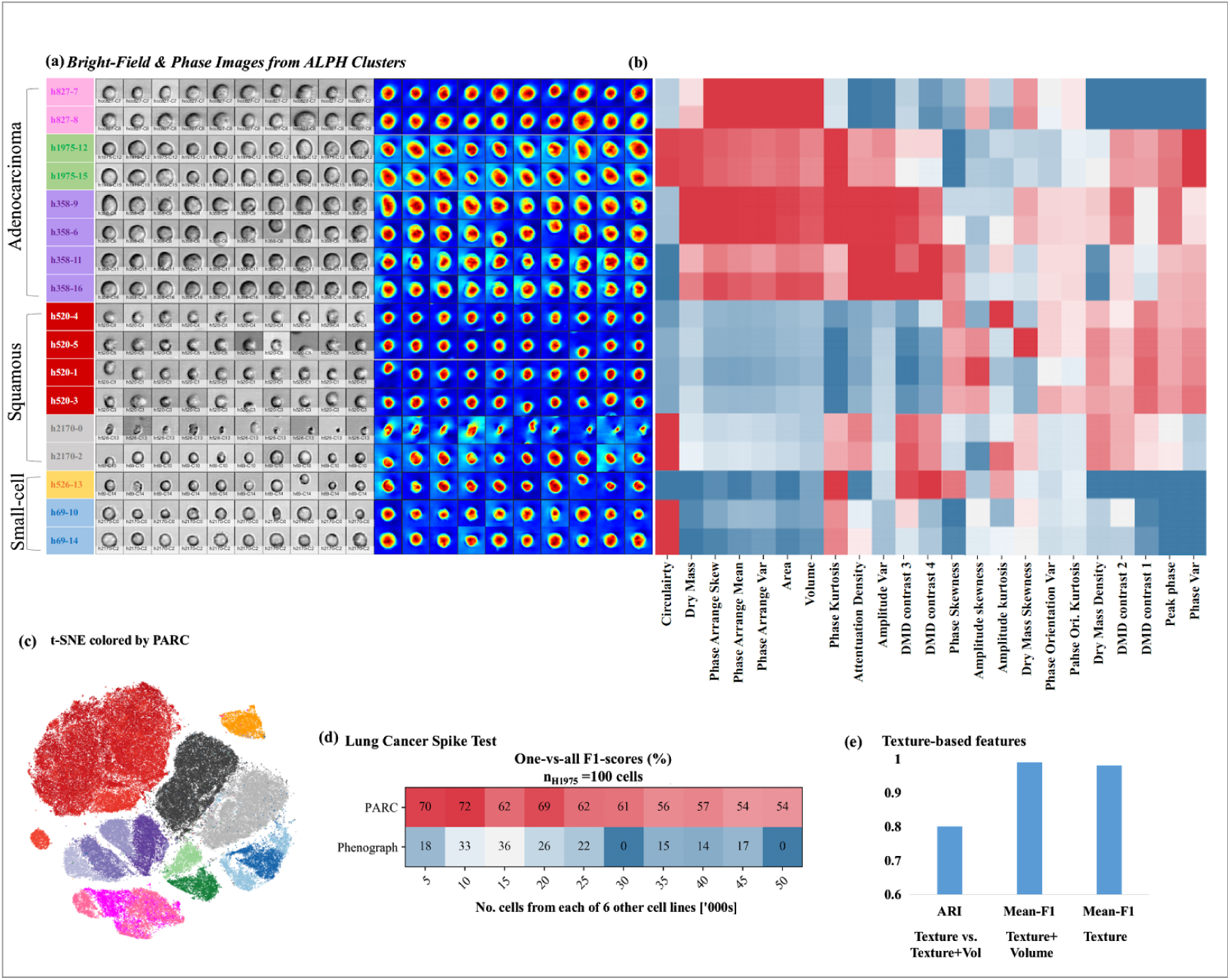
PARC for Ultralarge Analysis of Quantitative Label-free Single-Cell Image Data (1.1M lung cancer cells) by PARC. (a) (Left) Bright-field and (Right) quantitative phase images of cells captured by the multi-ATOM system. (b) Phenotypic profiles of the lung cancer cell populations clustered by PARC (See Suppl. Table 5 for the composition of the populations) based on a total 26 features related to the biophysical characteristics of single cells extracted from the multi-ATOM images (See Suppl. Table 6). Each of the three main lung cancer subtypes, squamous, adenocarcinoma, small-cell lung cancer, shows its characteristic phenotypic profile, with texture-based features further differentiating subtypes/clusters within the cell line. (d) A spike test where H1975 cells (n=100) are digitally mixed with progressively higher cell count of each of the other 6 lung cancer cell lines shows PARC’s ability to reliably detect the rare population. (e) PARC is employed to illustrate the significance of the label-free texture-based features of single-cells for distinguishing different cell types.

We further employ PARC to explore the scRNA-seq dataset of 1,308,421 embryonic mouse brain cells. The single-cell transcriptomic profiles were obtained with Cell Ranger 1.2 (10X Genomics 2017), and again preprocessed in the same manner as the Zheng et al 2017 dataset (see above) using python package SCANPY (Wolf et al 2018). Bypassing the approaches that downsample the data and thus the inevitable risk of losing the original data structure (especially the rare populations (Linderman 2019)), PARC completes the clustering with a run time of only 15 minutes on the 1.3 million cells (using the first 50PCs on the UMI counts of the 1000 most variable genes found after initial filtering). It is significantly faster than the competitive runtimes reported by the recent methods that do not rely on downsampling, i.e. ScScope (Deng 2019) and SCANPY (Wolf et al. 2018), with the clustering runtime of 97 and 104 minutes respectively. The clusters are annotated by the major cell types according to the maximal expression of well-known marker genes from the Allen Brain Atlas and Tasic et al 2016 (**Fig. 3c-f**), and have the following composition: GABAergic 18%, Glutamatergic 65% and non-neuronal 17%. The composition concurs with previous studies on *embryonic* brain cell composition which suggest ~90% of cells are neuronal (Bandeira et al. 2009), with ~1 in every 5 neurons being GABAergic (Sahara et al. 2012). The composition also agrees with the reported fractions by ScScope and SPLiT-Seq (Deng 2019) (**Fig. 3f**). Further classification of subtypes is inferred by plotting the average cluster expression for well-known gene markers, thus verifying the segregation of established (non-) neuronal types (**Fig. 3f** and **Supplementary Table 4**). Our results thus demonstrate the ability of PARC to enable efficient and effective exploration of the ultra-large heterogeneous single-cell datasets, which becomes increasingly demanding.

### 3.5 PARC clusters 1.1 million quantitative label-free single cell image data

An emerging challenge in single-cell analysis is to adapt to the progressively diverse sets of single-cell data generated by the wide range of new single-cell technologies, each with multiple modalities. This becomes a prerequisite for comprehensive multi-faceted single-cell analysis. Apart from the flow/mass cytometry and sequencing technologies, a notable example is high-throughput and high-content single-cell imaging - empowering large-scale deep image analysis that extracts a multitude of subtle features (or phenotypes) representing cell states and types (J. Caicedo et al. 2017).

In contrast to the fluorescence image cellular assay that specifically probes different biomolecular signatures of the cellular components and even provides functional annotation of genes by morphological similarity (Rohban et al 2017), a substantial body of work has shown that cellular biophysical properties, extracted from label-free optical imaging (Park et al. 2018; Zangle and Teitell. 2014; Tse et al. 2013; Otto et al. 2015), are the effective intrinsic markers for probing many cellular processes (e.g. cell proliferation, death, differentiation and malignancy). Bypassing the need for costly and time-consuming sample preparation, single-cell biophysical phenotyping could be functionally significant in single-cell analysis especially when other biomolecular assays are not effective.

Here we test the adaptability of PARC to cluster an in-house niche single-cell image dataset which describes the biophysical phenotypic profiles of 1.1 million lung cancer cells (7 cell lines representing three major lung-cancer subtypes: 1) adenocarcinoma, 2) squamous cell carcinoma, 3) small cell carcinoma). The biophysical phenotypes of individual cells were extracted from a recently developed ultrahigh-throughput microfluidic quantitative phase imaging (QPI) cytometer, coined multi-ATOM (Lee Feb 2019 and Lee April 2019), which captures a large amount of label-free single-cell images at an ultrahigh throughput (>10,000 cells/sec) without compromising sub-cellular resolution. In multi-ATOM, each imaged cell generates three different label-free image contrasts, from which 26 biophysical features are derived, e.g. cell size, mass, mass density, optical opacity, and statistical sub-cellular texture characteristics (See the detailed phenotype definitions in **Supplementary Table 5**). After feature normalization, we apply PARC to cluster the total 1,113,369 single cells imaged by multi-ATOM based on the 26 biophysical phenotypes.

PARC unambiguously separates (mean-F1 98.8%) between and within the 3 broad groups of lung cancer cells (**Fig. 4a-c**). Indeed, each of the three main lung cancer subtypes shows its characteristic phenotypic profile. We observe there are subtle differences in some texture features within the same subtype that further differentiate individual cell lines (**Fig. 4b**) - demonstrating the discriminative power of label-free biophysical phenotypes. PARC and Phenograph score the highest in terms of accuracy compared to the other methods (**Supplementary Fig. 1**), with PARC completing the task in 800 seconds versus the 7,200 seconds required by Phenograph using the same computational resources. Furthermore, by running PARC on the randomly selected n = 100 of H1975 cells mixed with an increasing cell count of each of the other six cell lines, we also demonstrate PARC’s consistent performance in rare-population detection based on biophysical phenotypes (**Fig. 4d and Supplemental Fig. 3**).

To exemplify its utility in image-based phenotypic exploration, we also use PARC to further investigate the significance of the label-free sub-cellular texture-based features in distinguishing different cell types. While cell size (volume) and shape are the most conceivable cellular biophysical features, sub-cellular textures parameterized from label-free imaging are intimately linked to a variety of subcellular spatial characteristics, e.g. protein localization (Yan et al., 2018), nucleus architectural changes (e.g. DNA fragmentation (Almassalha et al., 2016), and cytoskeletal network (Bon et al., 2014), to name a few. Hence, they can be harnessed as information-rich single-cell phenotypes. This can clearly be evident by the insignificant drop (1%) in the unweighted F1-score when exclusively the texture features (excluding volume, area, circularity and their moments) are input to PARC, compared to the case of using the complete feature set (**Fig. 4e**). The adjusted rand index (ARI) between the two sets of clusters of 80% indicates the two sets are well aligned.

## Discussion

The burgeoning of new bioassay technologies, notably the seamless integration of advanced molecular biology, microfluidics, and imaging, now allows diverse characterization of single cells at an unprecedented throughput and content. There is thus a pressing need for new computational tools that could efficiently handle such an increasing scale and complexity of single-cell data, and effectively explore the cellular heterogeneity for gaining new biological insights. To this end, PARC fills this gap by employing a combinatorial graph-based clustering approach that outperforms other competitive methods not only in speed (by an order of magnitude), and scalability (going beyond 1 million cells), but also the ability to accurately capture the complex data structure, especially to detect rare populations (< 0.1%).

PARC does not incur additional computational complexity to deal with exhaustive large-scale data processing or resort to random data downsampling. Instead, PARC is built upon three integrated elements to analyze ultralarge-scale and high-dimensional single-cell data: (1) HNSW for accelerated k-NN graph construction, (2) *data-driven* graph pruning and (3) a new community-detection approach, Leiden algorithm. Notably, our results show that pruning, guided by the local and global single-cell data structure, critically refines and improves the data graph representation that in turn further accelerates the Leiden algorithm, and alleviates the common problem of the resolution limit in community detection.

We anticipate that the clustering performance of PARC could readily be further augmented by incorporating other methodologies recently developed for single-cell analysis. For instance, prior to PARC clustering, one could apply correction steps for removing the batch effects (MNN by Haghverdi et *al., 2018*), imputation strategies for combating the noise and dropouts in scRNA-seq data (e.g. scScope, DeepImpute) and accelerated visualization methods for ultralarge-scale single-cell data (e.g. Net-SNE, flt-SNE and UMAP). Such compatibility, together with the fact that PARC does not require prior knowledge of the single-cell data, could make PARC widely adaptable to the popular single-cell analysis pipelines (e.g. Seurat, SCANPY and Cell Ranger) or new methodologies tailored for niche single-cell data types (e.g. label-free imaging cytometry data shown in **Fig. 4**) and integration of multiple data types.

Indeed, our results demonstrate that PARC produces robust and accurate clustering across various single-cell data types, namely flow cytometry, mass cytometry, scRNA-seq and even emerging imaging cytometry. We thus anticipate that such versatility of PARC could be extended to play an important role in emerging techniques that empower multi-faceted and integrative characterization of single-cell biochemical/biophysical phenotypes and transcriptional profiles (or broadly regarded as single-cell multi-omics (Hasin et al. 2017, Chappell et al. 2018) – the major pursuit to crafting the human cell atlas (Regev et al., 2017) that could offer a deeper mechanistic understanding of biological processes, particularly those driving cellular heterogeneity associated with diseases.

## Declaration of Interests

The authors declare no competing interests.

## Notes

https://github.com/ShobiStassen/PARC

