## Supplementary Information for "PARC: ultrafast and accurate clustering of phenotypic data of millions of single cells"

### Supplementary Data

#### LEAD CONTACT AND MATERIALS AVAILABILITY

This study did not generate new unique reagents

#### QUANTIFICATION AND STATISTICAL ANALYSIS

In this section we describe the choice of performance measures used. For the multi-population mass cytometry datasets where relatively granular annotations where available, we adopt the approach used by Weber and Robinson 2016 and Samusik et al 2016 to compute the F1-measure as follows: First construct an F1-matrix and then apply the Hungarian algorithm to find an optimal one-to-one assignment between the manually-gated populations and the automatically detected clusters. $F_{ij}=2(R_{ij}P_{ij})/(R_{ij}+P_{ij})$ is the ij'th entry of the F1-matrix. Precision $P_{ij}$ is the number of matches between a true class and a cluster, divided by the number of cells in that cluster; and Recall $R_{ij}$ as the number of matches between the true label and the cluster, divided by the total number of cells for that true label (that should have been in that cluster). Precision and Recall are defined as $P_{ij} = C_{ij}/\Sigma_j(C_{ij})$, and $R_{ij} = C_{ij}/\Sigma_k(C_{kj})$ respectively. $Cij$ is the number of cells in the i'th cluster that belong to the j'th reference population [45].

The method offers some advantages over the more common multi-class extension of the F1-score (where $T$ is the set of true classes, $T = \{t_1, t_2, ..., t_n\}$ and $S$ is the clustering result $= \{s_1, s_2, ..., s_m\}$.

$$F(T,S)=\sum_{t_i \in T}\frac{t_i}{N}\max_{s_j \in S}\left\{F\left(t_i,s_j\right)\right\}$$

Most evidently, it does not bias the overall F1-measure to that found in larger populations (thus obscuring the importance of smaller but phenotypically distinct populations) and less trivially, it also avoids the problem of potentially assigning multiple reference populations to the same cluster.

However, in the absence of an adequately well annotated or detailed 'ground truth', the Hungarian method can be punitive when coarse manual gatings overlook real divisions (subpopulations) revealed by the clusters.

For the 10X PBMC scRNA-seq data as well as the Multi-ATOM data, the 'ground truth' reference annotations are coarse. To avoid penalizing the algorithm for splitting up an annotated population into distinct subtypes (e.g. a reference monocyte annotation may have been divided into clusters we can infer as classical and non-classical monocytes), we evaluate the performance evaluation based on a macro F1-score. We assign each cluster a reference label based on its majority population. This means there may be 1 or more clusters that are assigned to a particular reference population and belong to that 'macro-level' cluster. We then compute the one-vs-all F1-score for each of the major annotated populations on the 'macro-cluster' level. The mean one-vs-all F1-scores are an unweighted average to give importance to rare cells. When using the macro-F1-score, the number of clusters uncovered is reported in order to highlight that performance is achieved within a number of clusters suitable for downstream analysis.

### DATA AND CODE AVAILABILITY

PARC source code is available from github: https://github.com/ShobiStassen/PARC

The mass cytometry and flow cytometry datasets (Mosmann_rare, Nilsson_rare, Samusik_all, Levine_32dim, Levine_13dim) are all publicly available at FlowRepository (repository I.D.: FR-FCM-ZZPH)

The Mouse Brain and Zheng_PBMC sc-RNA datasets are publicly available from 10X Genomics website https://www.10xgenomics.com/solutions/single-cell/

The Multi_ATOM lung cancer data supporting the current study are available from Mendeley https://data.mendeley.com/datasets/nnbfwjvmvw/draft?a=dae895d4-25cd-4bdf-b3e4-57dd31c11e37

### EXPERIMENTAL MODEL AND SUBJECT DETAILS OF LUNG CANCER IMAGING FLOW CYTOMETRY DATASET

##### Multi-ATOM imaging protocol for Lung Cancer Data

Multi-ATOM is an ultrafast quantitative phase imaging (QPI) technique that bypasses the use of camera technology and its speed limitation. It measures the optical path length of the cell for deriving multiple biophysical markers, e.g. cell size, morphology, mass density, sub-cellular texture, refractive index etc. (Park et al. 2018, and Lee April 2019). Detailed configuration of multi-ATOM can be referred to (K. Lee et al., Feb 2019 and K. Lee et al., April 2019). In brief, it relies on all-optical image encoding in and retrieval from the broadband laser pulses by two mapping steps at an ultrafast line-scan rate governed by the laser repetition rate (11.8 MHz in our case): wavelength–time mapping (time-stretch process) and wavelength–space mapping (spectral-encoding process). Here the cells are in a unidirectional microfluidic flow orthogonal to the spectrally-encoded line-illumination, at a flow speed >1 m/s (equivalent to a typical imaging throughput of 10,000 cells/s). These encoded line-scans are then digitally stacked to form the two-dimensional (2-D) images of the cells. Furthermore, by means of multiplexed spectral-encoding measurements, multi-ATOM retrieves the spatially dependent optical phase shift, or simply called *quantitative phase* from each line-scan – a quantity closely linked to variations in cell morphology and refractive index distribution that cause optical wavefront distortion as the illumination light propagates through a cell. Following our previous work (K. Lee et al., Feb 2019), we ensured robust in-focus single-cell imaging under a fast-microfluidic flow (>1 m/s) by designing the microfluidic channel platform to optimize the balance between the inertial lift force and the viscous drag force. The microfluidic channel was fabricated by curing polydimethylsiloxane (PDMS) on a silicon wafer mold which was prepared by the standard soft lithography technique.

The enormous number of image-context-rich multi-ATOM images favours analysis of the high-dimensional *spatially-resolved* biophysical single-cell data in large-scale. Based on the amplitude and quantitative phase images of cells, we extract the typical bulk parameters, e.g. size, averaged dry mass density (DMD), and optical opacity. Beyond that, we further transform the quantitative phase image into a 2D *dry-mass-density contrast (DC) map* and analyse its spatial variation in a statistical histogram (**Supplementary Table 5**). We note that the DC map visualizes the *local* variation of DMD within the cell. Based on the amplitude, quantitative phase and DC images, we extract in total **26** features, representing different aspects of biophysical properties of single cells.

**Cell-culture of lung cancer cell lines**

The 7 lung cancer cell lines consisted of 5 adherent (H358, H1975, HCC827, H520 and H2170) and 2 suspended (H526 and H69) lines. They were all cultured in their full medium: Roswell Park Memorial Institute (RPMI)-1640 with ATCC modification (Gibco™), supplemented with 10% Fetal Bovine Serum (FBS, GGibco™) and 1% Antibiotic-Antimycotic (Gibco™). The adherent types were trypsinized by 0.25% Trypsin-EDTA (Gibco™) for 4 minutes at 37ºC. The detached cells were extracted, centrifuged and re-suspended in complete medium. A portion would be re-seeded and the remaining (usually $1 \times 10^6 \sim 3 \times 10^6$) suspended in 3mL of complete medium prior to commencing the flow experiment. The suspended cell lines were centrifuged and re-suspended in complete medium. The same strategy was used to split a portion for re-seeding and the remaining were suspended in 3mL complete medium for the flow experiment. The experiment was held within 3 hours of harvesting.

### KEY RESOURCES TABLE

| REAGENT or RESOURCE | SOURCE | IDENTIFIER |
|---|---|---|
| Software and algorithms | | |
| PARC | this paper | https://github.com/ShobiStassen/PARC-phenotyping-by-accelerated-refined-community-partitioning.git |
| Hierarchical Navigable Small World | Malkov and Yashunin, 2016 | arXiv: 1603.09320 and https://github.com/nmslib/hnswlib |
| igraph | python package | https://igraph.org/python/ |
| multicore tsne | | https://github.com/DmitryUlyanov/Multicore-TSNE |
| Scanpy | Wolf et al. 2018 | https://scanpy.readthedocs.io/en/stable/ |
| Leiden algorithm | Traag et al. 2019 | https://github.com/vtraag/leidenalg.git |
| Deposited data | | |
| Samusik_all | Samusik et al (2016) | doi.org/10.1038/nmeth.3863 and doi.org/10.1002/cyto.a.23030 |
| Levine_32dim and Levine 13dim | Levine et al (2015) | doi: 10.1016/j.cell.2015.05.047 and 10.1002/cyto.a.23030 |
| Mosmann_rare | Mosmann et al (2014) and Weber and Robinson 2016 | doi.org/10.1002/cyto.a.22445 and doi.org/10.1002/cyto.a.23030 |
| Nilsson_rare | Nilsson et al 2013 and Weber and Robinson 2016 | doi.org/10.1002/cyto.a.23030 and doi.org/10.1002/cyto.a.23030 |
| 10X PBMC | Zheng et al 2017 | 10.1038/ncomms14049 and www.10xgenomics.com/solutions/single-cell/ |
| 10X 1.3Million Mouse Brain | 10X Genomics 2017 | www.10xgenomics.com/solutions/single-cell/ |

### Supplementary Figures

#### Supplementary Figure 1

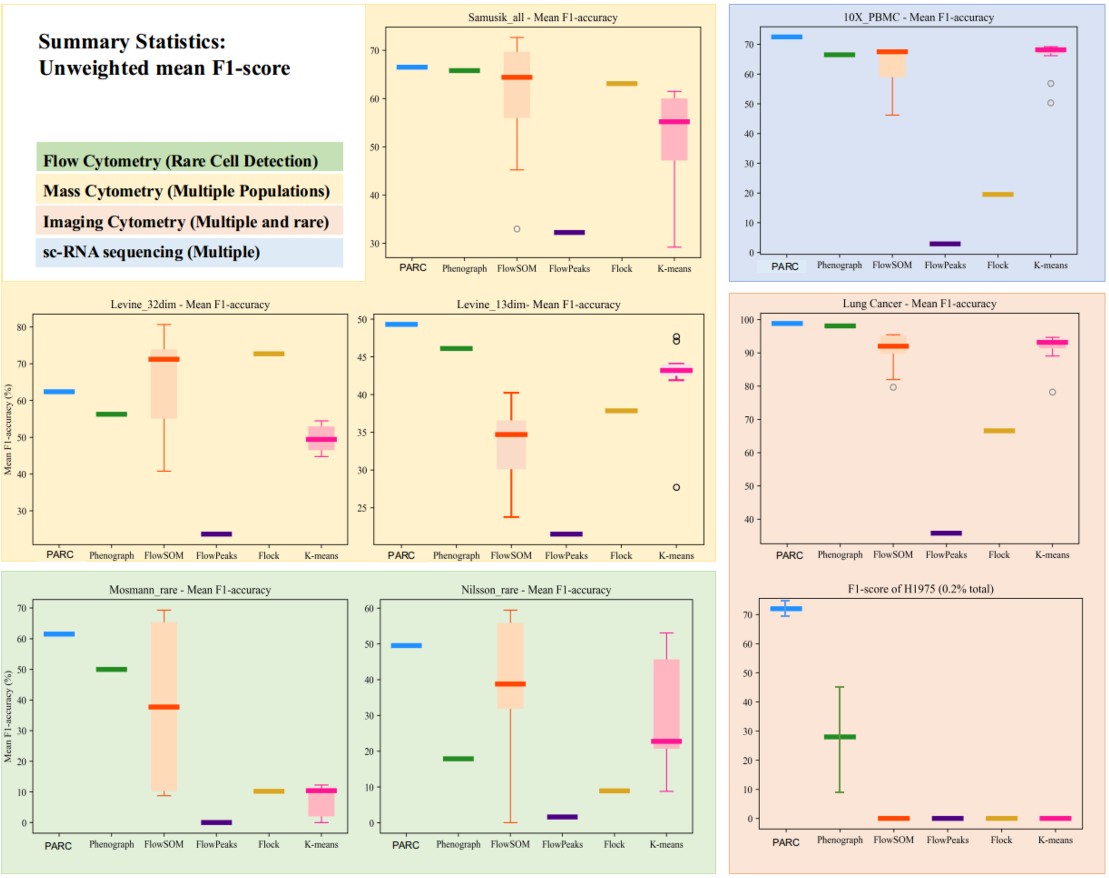

### Supplementary Figure 2

#### Multiple population detection mean F1-Score and No. of Clusters

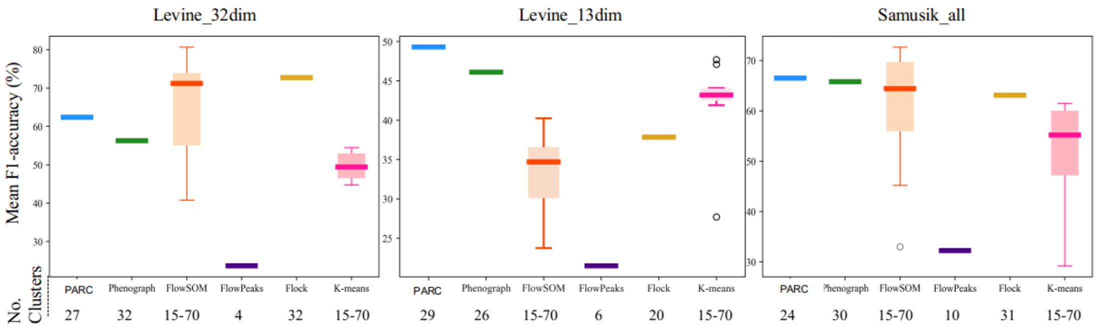


### Supplementary Figure 3

#### Stability of Rare-cell detection of 100 H1975 cells: Phenograph vs. PARC

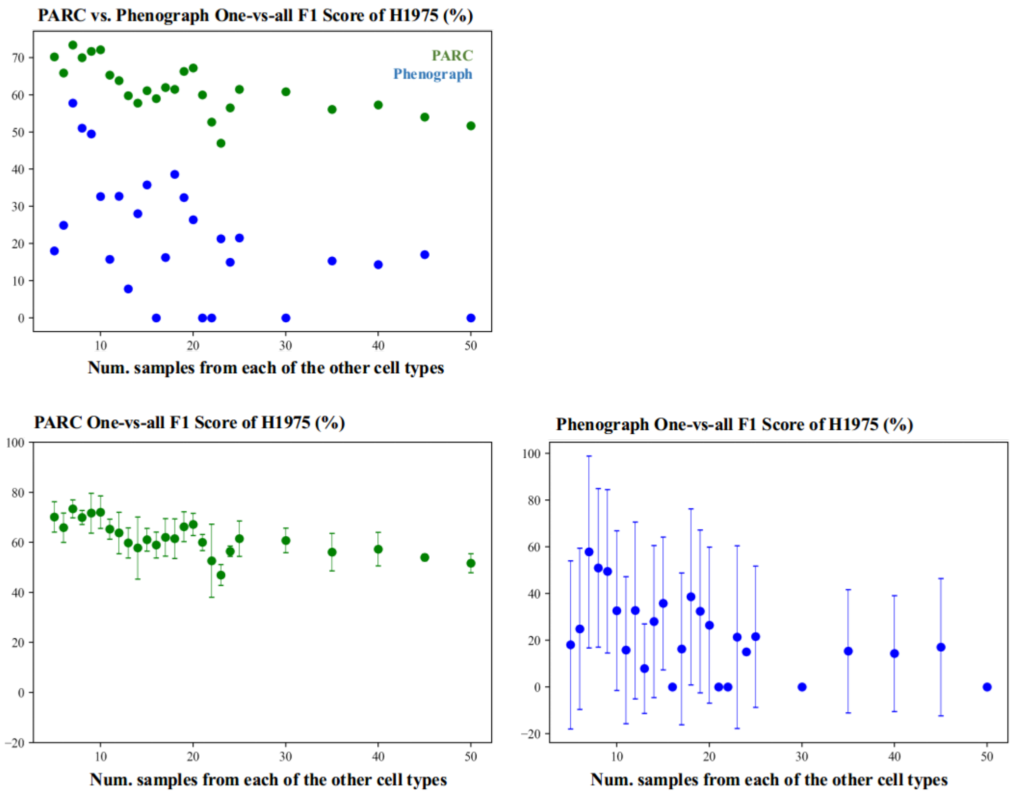

### Supplementary Figure 4

##### "Ground Truth" annotations of 68K 10X PBMC sc-RNA data

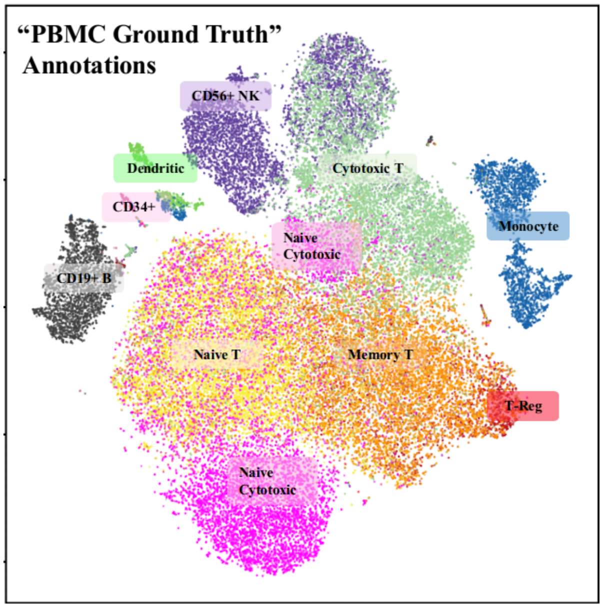


### Supplementary Table 1

##### Performance characteristics: PARC avoids excessive fragmentation and offers significant speedup

| *Multi-pop* | **Levine_32dim** N = 265K | | | **Levine_13dim** N = 167K | | | **Samusik_All** N = 841K | | |
|---|---|---|---|---|---|---|---|---|---|
| | **F1** | **clus** | **time (s)** | **F1** | **clus** | **time (s)** | **F1** | **clus** | **time (s)** |
| *PARC* | 0.63 | 28 | 90 | 0.49 | 25 | 35 | 0.66 | 24 | 331 |
| *Pheno* | 0.56 | 32 | 1537 | 0.46 | 27 | 984 | 0.65 | 30 | 8549 |
| *X-shift* | 0.69 | 31 | 11125 | 0.47 | 153 | 2897 | 0.66 | 74 | 13700 |

### Supplementary Table 2

**Pruning ensures rare populations to be consistently captured, whereas lowering K only marginally improves rare cell detection at the cost of fragmentation**

| *Rare* | Nilsson_rare N= 44K | | Mosmann_rare N= 396K | | H1975 (0.04%) N= 280,100 | |
|---|---|---|---|---|---|---|
| **Phenograph** | **F1** | **clus** | **F1** | **clus** | **F1** | **clus** |
| ***K=10*** | 0.21 | 39 | 0.15 | 36 | 0 | 35 |
| ***K=15*** | 0.37 | 33 | 0.46 | 31 | 0 | 27 |
| ***K=20*** | 0.18 | 30 | 0.49 | 24 | 0 | 22 |
| ***K=25*** | 0.19 | 28 | 0.48 | 24 | 0 | 20 |
| ***K=30 (default)*** | 0.18 | 26 | 0.50 | 20 | 0 | 18 |
| ***PARC*** | | | | | | |
| ***K=30 default pruning*** | **0.49** | **31** | **0.62** | **20** | **0.55** | **24** |
| ***K=30, no pruning*** | 0.14 | 19 | 0.00 | 14 | 0 | 14 |
| ***K=10, no pruning*** | 0.18 | 23 | 0.66 | 19 | 0.24 | 24 |
| ***K=15, no pruning*** | 0.18 | 19 | 0.60 | 19 | 0 | 18 |
| ***K=20, no pruning*** | 0.15 | 19 | 0.01 | 16 | 0 | 19 |
| ***K=25, no pruning*** | 0.14 | 18 | 0.01 | 16 | 0 | 16 |

### Supplementary Table 3

##### 10X PBMC (68,000 cells) Marker genes and references

| Markers for cell type inference | | | |
|---|---|---|---|
| **Cluster** | **Inferred Cell type** | **Markers** | **[Various Ref]** |
| 9 | Classical Monocyte | CD14+ FCGR3A- (CD16-), CX3CR1-, S100A12+ | Ajami and Steinman 2018: Wong et al. (2011) ,Schinnerling et al. 2015; Stansfield and Ingram (2015) |
| 10 | Non-Classical Monocyte | CD16+, CD14+, CX3CR1+, S100A12- | |
| 13 | Myeloid and Monocyte related CD14+ Dendritic | CD1C+, CD1E+, HLA-genes, CD14 | Collin et al. (2013); Wojciech (2011) |
| 11 | Plasmacytoid Dendritic | IL-3RA+ (CD123), GZMB | Collin et al. (2013); Tel et al. (2011), Zhang et al., 2017 |
| 5 | NK cytotoxic CD56dimCD16+ | NKG7+, FCGR3A+(CD16+), CD160+, CCR7- | Le Bouteiller et al. (2011), Hong (2012), Turman (1993) |
| 3 | NK II CD56brightCD16dim | CD160-, NKG7+ | Le Bouteiller et al. (2011), Turman (1993) |
| 8 | Mature B cell | CD22+, CD79A+, CD79B+, CD19+ | Boyd (2016) |
| 17 | Early-B cell | IGJ+, CD79A+ | Hystad (2007), Boyd (2016) |
| 12 | CD34+Megakaryocyte | PF4+,GP9+, TREML1+, PPBP+ | Sakurai (2016), Smith (2018) |
| 6 | CD25+ T-Reg | FOXP3+, CD52+, CD70+ | Rudensky (2011), Samten (2013) |
| 2 | CD8+ Naive Cytoxic | CCR7+ CD8A+, CD4- CD28-, GZMK | Campbell (2001) |
| 4,7 | CD8+ CytoToxic (activated) | CD8A+, NKG7 | Turman (1993) |
| 1 | CD4+ T-Memory | CD8A-, CCR7- | Campbell (2001) |
| 16 | CMV-specific CD4+ T-Memory | IFI6+, IFI27+, IFIT3 | Hu (2013) |
| 0,15,14 | CD4+ Naive T | CD4+, CD8A-, CCR7+ | Campbell (2001) |

### Supplementary Table 4

**Marker Genes and references for 10X Mouse Brain (1.3 Million cells)**

| Cell sub-types | References |
|---|---|
| **Tbr1, Eomes, Slc16a7, Slc16a7** | Zeisel et al., 2018 |
| Thalamic origin (Slc16a7) | Liguz-Lecznar and Skangiel-Kramska 2017 |
| pyramidal projection neurons (Eomes, Tbr1) | Hevner et at., 2006 |
| Cortical Terminals (Slc16a6) | Liguz-Lecznar and Skangiel-Kramska 2017 |
| **Aldoc, Hes1, Olig1, Gja1, Gfap** | |
| Aldoc, Gfap (astrocyte) | Tasic et al., 2016<br>Boisvert et al., 2018 |
| Olig1 (oligdendrocyte) | Othman te al., 2011 |
| Hes1 (Glia) | Furukawa et al., 2000 |
| **Gad1, Gad2, Slc32a1** | Tasic et al., 2016 |
| Sst (SOM interneuron) | Gonchar et al., 2007 |
| Calcr (CR interneuron) | Reynolds and Beasley 2001 |
| Calb1 (CB interneuron) | Reynolds and Beasley 2001 |
| Htr3a (cortical interneuron) | Frazer et al., 2017 |

### Supplementary Table 5

#### Population composition of PARC's clustering of Lung Cancer multi-ATOM data

| Cell type | Major cell line | Cell count |
|---|---|---|
| h2170-0 | squamous | 167662 |
| h2170-2 | squamous | 129825 |
| h526-13 | small cell | 31058 |
| h520-1 | squamous | 154626 |
| h520-3 | squamous | 126852 |
| h520-4 | squamous | 60130 |
| h520-5 | squamous | 57626 |
| h358-6 | Adenocarcinoma | 53719 |
| h358-9 | Adenocarcinoma | 39385 |
| h358-11 | Adenocarcinoma | 33653 |
| h358-16 | Adenocarcinoma | 22751 |
| h69-10 | small cell | 35884 |
| h69-14 | small cell | 28802 |
| h827-7 | Adenocarcinoma | 47259 |
| h827-8 | Adenocarcinoma | 45029 |
| h1975-12 | Adenocarcinoma | 30069 |
| h1975-15 | Adenocarcinoma | 22648 |

### Supplementary Table 6

#### Description of biophysical features extracted from multi-ATOM images

**Biophysical features summary**

| Feature | QPI/BF | Description |
|---|---|---|
| Area/Volume | BF | Linked to cell proliferation and growth and often used in conjunction with cell mass to study regulation of cell size (Girshovitz and Shaked 2012) |
| Circularity | BF | Measure of conic deviation from cell being circular. Can be indicative of cell apoptosis or disease mass to study regulation of cell size (Girshovitz and Shaked 2012) |
| Attenuation Density | BF | Indicative of cell composition |
| Amplitude (Moments) | BF | Peak, mean and variance of BF amplitude |
| Dry Mass Density | QPI | Mass of non-aqueous material of cell, which is the integral of the optical path delay profile on the projected cell area. mass to study regulation of cell size (Girshovitz and Shaked 2012) |
| Dry Mass Density Contrast (Moments) | QPI | Statistical features of DMD contrast which measures local variations in DMD (Lee 2018) |
| Phase Arrangement (Moments) | QPI | Extract spatial phase information by considering the distribution of the phase as a function of the radius (distance from image center) |
| Phase orientation | QPI | Distribution of the phase as a function of a product of the angle and radius (from center of image) |

BF: bright-field; QPI: quantitative phase image; DMD: dry-mass density
